# Involvement of ArlI, ArlJ, and CirA in Archaeal Type-IV Pilin-Mediated Motility Regulation

**DOI:** 10.1101/2024.03.04.583388

**Authors:** Priyanka Chatterjee, Marco A. Garcia, Jacob A. Cote, Kun Yun, Georgio P. Legerme, Rumi Habib, Manuela Tripepi, Criston Young, Daniel Kulp, Mike Dyall-Smith, Mecky Pohlschroder

## Abstract

Many prokaryotes use swimming motility to move toward favorable conditions and escape adverse surroundings. Regulatory mechanisms governing bacterial flagella-driven motility are well-established, however, little is yet known about the regulation underlying swimming motility propelled by the archaeal cell surface structure, the archaella. Previous research showed that deletion of the adhesion pilins (PilA1-6), subunits of the type IV pili cell surface structure, renders the model archaeon *Haloferax volcanii* non-motile. In this study, we used EMS mutagenesis and a motility assay to identify motile suppressors of the Δ*pilA*[*1-6*] strain. Of the eight suppressors identified, six contain missense mutations in archaella biosynthesis genes, *arlI* and *arlJ*. Overexpression of these *arlI* and *arlJ* mutant constructs in the respective multi-deletion strains Δ*pilA*[*1-6*]Δ*arlI* and Δ*pilA*[*1-6*]Δ*arlJ* confirmed their role in suppressing the Δ*pilA*[*1-6*] motility defect. Additionally, three suppressors harbor co-occurring disruptive missense and nonsense mutations in *cirA*, a gene encoding a proposed regulatory protein. A deletion of *cirA* resulted in hypermotility, while *cirA* overexpression in wild-type cells led to decreased motility. Moreover, qRT-PCR analysis revealed that in wild-type cells, higher expression levels of *arlI*, *arlJ*, and the archaellin gene *arlA1* were observed in motile early-log phase rod-shaped cells compared to non-motile mid-log phase disk-shaped cells. Conversely, Δ*cirA* cells, which form rods during both early and mid-log phases, exhibited similar expression levels of *arl* genes in both growth phases. Our findings contribute to a deeper understanding of the mechanisms governing archaeal motility, highlighting the involvement of ArlI, ArlJ, and CirA in pilin-mediated motility regulation.

**Importance:** Archaea are close relatives of eukaryotes and play crucial ecological roles. Certain behaviors, such as swimming motility, are thought to be important for archaeal environmental adaptation. Archaella, the archaeal motility appendages, are evolutionarily distinct from bacterial flagella, and the regulatory mechanisms driving archaeal motility are largely unknown. Previous research has linked the loss of type IV pili subunits to archaeal motility suppression. This study reveals three *Haloferax volcanii* proteins involved in pilin-mediated motility regulation, offering a deeper understanding of motility regulation in this understudied domain while also paving the way for uncovering novel mechanisms that govern archaeal motility. Understanding archaeal cellular processes will help elucidate the ecological roles of archaea as well as the evolution of these processes across domains.

## Introduction

Microorganisms have evolved sophisticated motility machineries that allow them to adeptly navigate their diverse environments, for example to acquire nutrients or to avoid toxins^1,2^. In Bacteria, the rotation of flagella promotes swimming motility. However, in Archaea, evolutionarily distinct rotating surface filaments called archaella drive swimming motility^3–5^. The biosynthesis machinery of archaella is evolutionarily related to that of type IV pili (T4P), appendages required for bacterial and archaeal surface motility and adhesive interactions required for biofilm formation^6–10^. For instance, the subunits of both structures—archaellins in archaella and pilins in T4P—share class III signal peptides, which are required for transport to the cell surface and are processed by homologous signal peptidases^8^. Moreover, key assembly proteins, such as ArlI and ArlJ for archaella, and PilB and PilC for type IV pili, also exhibit remarkable homology, hinting at a deeper evolutionary connection between these seemingly disparate cell surface structures.

The archaellin genes are often found in a gene cluster^11–13^ along with genes encoding components of the archaellum motor complex, such as the ATPase ArlI, which assembles into the hexameric rotor, and the membrane protein anchor ArlJ^11,13^. The components of the archaellum motor also include the ATP-binding protein ArlH^14^ and stator proteins ArlF and ArlG^15,16^. In Euryarchaeota (Methanobacteriota^17^), ArlC/D/E, or any combination or fusion of the three, are thought to serve as a scaffold for the motor^7,18^. In Crenarchaeota (Thermoproteota^19^), a putative membrane protein ArlX is thought to replace ArlC/D/E^13,15^.

Recent studies have elucidated the regulatory roles of proteins encoded by genes in the *arl* cluster of some archaea. Both the euryarchaeal *Pyrococcus furiosus* and the crenarchaeal *Sulfolobus acidocaldarius* ArlH homologs have been shown to autophosphorylate. In the non-phosphorylated state, ArlH hexamerizes and associates with the ArlI rotor, but upon phosphorylation of ArlH, its affinity for ArlI is reduced. Deleting *arlH* or abrogating the phosphorylation of ArlH results in a loss of cell motility, signifying an important regulatory role for ArlH^14,20,21^. While little is known about other aspects of archaella regulation, many euryarchaeal ArlH-homologs contain a KaiC domain. KaiC family ATPases, which are named after a key component of the circadian clock proteins of cyanobacteria, have been proposed to have signal transduction roles in archaea^22^.

The motility phenotype of the euryarchaeon, *Haloferax volcanii*, correlates with its cell shape. Transitions between motile rod-shaped cells and sessile disk-shaped cells can be observed in liquid culture^23^, or in motility halos formed after cells are stab-inoculated into soft agar^24^. Cells observed in early-log growth phase or at the edge of the motility halos are motile, rod- shaped, and exhibit archaella on their surface. In mid-to-late log growth phase or at the center of the motility halos, they present as sessile, non-archaellated, pleomorphic disks^23,24^. Reported mutants that cannot form rods are unable to swim^24,25^. Conversely, mutants that are locked in the rod shape tend to be hypermotile^25^.

In our previous study, two mutants isolated from a hypermotility screen were rod-only in the conditions tested. Those mutants had premature stop codons in the gene *cirA* which encodes Circadian regulator CirA^25^. Similar to ArlH, CirA is also a KaiC-like protein, and *cirA* is located within the *arl* gene cluster^22^. Like the cyanobacterial circadian clock proteins, transcription of *cirA* oscillates with a light/dark cycle^26^. Thus, the positioning of *cirA* proximal to the *arl* genes, along with the proposed importance of KaiC-like proteins in archaeal signal transduction^22^, suggests that CirA may play a role in motility regulation.

Esquivel *et. al* reported that the conserved hydrophobic domain (H-domain) of type IV pilins is not only required for the adhesion, but is also involved in motility regulation^27^. This N-terminal H-domain is critical for the assembly of pilins into a pilus fiber^6^ and is also part of the class III signal peptide. This signal peptide contains a unique signal peptidase cleavage site N-terminal to the H-domain^28^. Upon cleavage, processed pilins are incorporated into a pilus. It has been proposed that a subset of processed pilins is retained in the membranes of planktonic cells. Esquivel *et al*. found that deleting all six *Hfx. volcanii* adhesion pilins containing this conserved H-domain (Δ*pilA*[*1-6*]) results in a loss of both adhesion and swimming motility. Expression *in trans* of the H-domain fused to a non-pilin protein is unable to restore surface adhesion, but can restore motility in these cells^27^. This suggests that, in addition to increasing surface adhesion, the assembly of these pilins into pili may also promote biofilm formation by preventing membrane- associated pilins from enabling archaella-driven motility. The mechanism by which the depletion of pilins inhibits motility is not well-understood.

Here, we describe a motility screen that identified suppressor mutants of the non-motile Δ*pilA*[*1-6*]. Most of the suppressors harbor missense point mutations in *arlI* and/or *arlJ*. Using reverse genetics, we have demonstrated that these mutations enable cells to swim despite lacking the pilin H-domains. Additionally, a subset of these suppressors harbors disruptive mutations in *cirA*. Transcriptional analysis of a Δ*cirA* strain suggests that CirA reduces expression of the *arl* genes, revealing a novel avenue of motility regulation.

## Results

### Ethyl methanesulfonate (EMS) mutagenesis of **Δ***pilA*[*1-6*] revealed motile suppressors

To identify components involved in H-domain dependent motility regulation, we began by screening for suppressors of the motility defect of the Δ*pilA*[*1-6*] mutant strain. Since spontaneous motile mutants of Δ*pilA*[*1-6*] streaked across a soft-agar motility plate were not identified after incubation for 5 days, we screened with chemically induced mutations. Cells were incubated with 0.05 M EMS for 0, 10, 20, and 30 minutes. The 20-minute incubation was the shortest exposure time that resulted in suppressors that could form motility halos (**Fig. 1**). After streaking cells from the edge of the motility halo onto solid agar plates, isolated colonies were picked and stabbed onto soft agar motility plates to confirm the motility phenotype. Eight suppressors of the Δ*pilA*[*1-6*] motility defect were isolated using this method.

**Figure 1.**
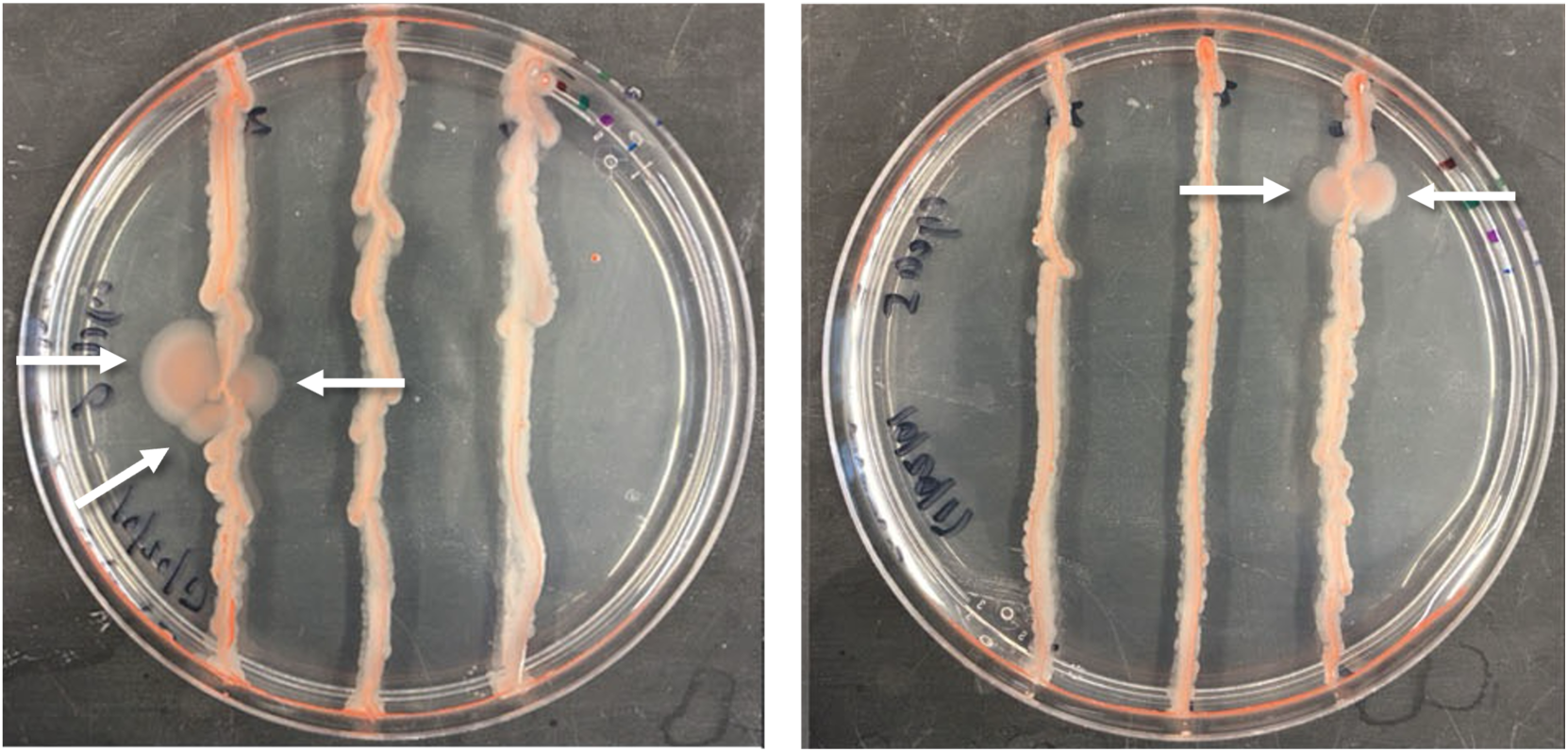
EMS mutagenesis results in Δ*pilA*[*1-6*] suppressor mutants that regain swimming motility. Two representative images of soft agar plates (0.3% w/v) that have been streaked with *Hfx. volcanii* Δ*pilA*[*1-6*] incubated with 0.05 M EMS for 20 minutes. White arrows point to outgrowths of motile suppressors after five days of incubation at 45°C.

### Majority of Δ*pilA*[*1-6*] suppressor mutants contain mutations in *arlI, arlJ*, and/or *cirA*

To identify the genes altered by EMS mutagenesis, genomic DNA isolated from the eight suppressor mutants was sent for whole genome sequencing (WGS) (**Table S1**). All of the suppressor strains had multiple mutations. The total number of mutations ranged between 7 and 35, not including the engineered deletion of PilA[1-6]. Most mutations occur in only one suppressor strain. Only three genes (*arlI*, *arlJ*, and *cirA*) were identified as having distinct mutations in at least two Δ*pilA*[*1-6*] suppressor strains (**Table 1, Table S1**), suggesting the significance of these genes in the suppressor phenotype. Moreover, *arlI*, *arlJ*, and *cirA* are all located in the same genomic region, strongly suggesting a link to pilin-mediated motility regulation. While multiple mutants encoding an ArlI^A331V^ and ArlJ^A364V^ were identified, these strains have different secondary mutations (**Table S1**) and total number of mutations, suggesting that the mutations in these specific residues may play a role in the suppression of the motility defect of Δ*pilA*[*1-6*]. Two suppressors, KY-7 and KY-8, have mutations in both *arlI* and *cirA*, and one suppressor, KY-1, includes mutations in all three genes (**Table 1**). This suggests that the proteins encoded by these genes may work in combination to regulate motility. ArlJ^A364V^ does not co-occur with mutations in either ArlI or CirA, which may indicate a significant role of this residue in motility regulation.

**Table 1.**
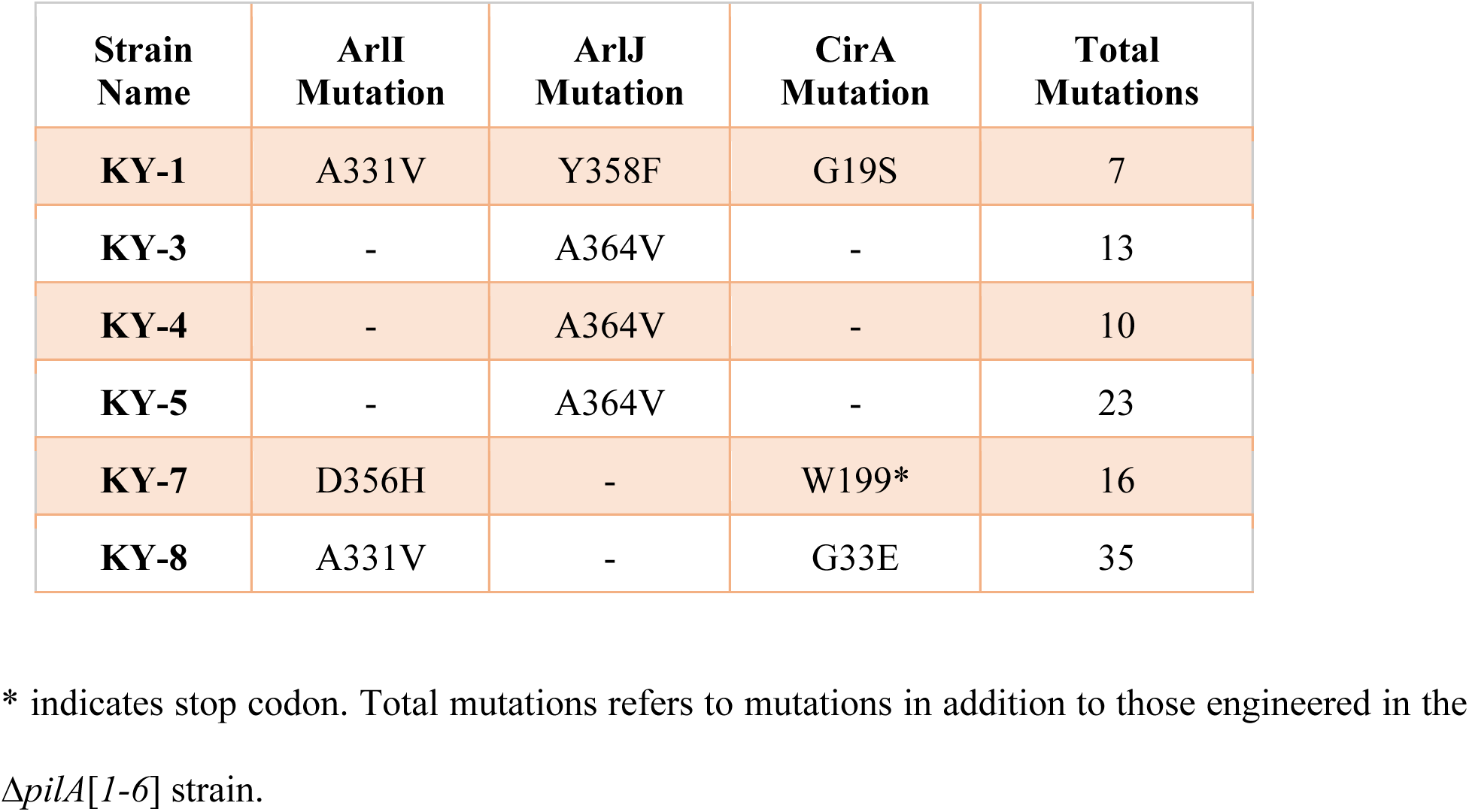

| **Strain Name** | **ArlI Mutation** | **ArlJ Mutation** | **CirA Mutation** | **Total Mutations** |
| --- | --- | --- | --- | --- |
| **KY-1** | A331V | Y358F | G19S | 7 |
| **KY-3** | - | A364V | - | 13 |
| **KY-4** | - | A364V | - | 10 |
| **KY-5** | - | A364V | - | 23 |
| **KY-7** | D356H | - | W199* | 16 |
| **KY-8** | A331V | - | G33E | 35 |

\* indicates stop codon. Total mutations refers to mutations in addition to those engineered in the *ΔpilA[1-6]* strain.

### Δ*pilA*[*1-6*] suppressors have mutations that may slightly alter the functions of ArlI and ArlJ but disrupt CirA

In the Δ*pilA*[*1-6*] motile suppressors, most of the amino acid substitutions in ArlI and ArlJ retain the same chemical properties as the replaced residue, such as in the alanine or glycine substitutions for valine in ArlI^A331V^ and ArlJ^A364V^, which retain aliphaticity, and the tyrosine substitution for phenylalanine in ArlJ^Y358F^, which retains aromaticity. ArlI^D356H^ is the only mutation in ArlI and ArlJ that results in a major change in the properties of the residue: aspartate changed to histidine (**Table 1**). To assess whether the mutated residues affect conserved regions of the proteins, we aligned the *Hfx. volcanii* ArlI and ArlJ protein sequences with those of four haloarchaeal homologs, as well as five homologs from representative archaeal species of different phyla (**Fig. 2a, 2b, Fig. S1**).

**Figure 2.**
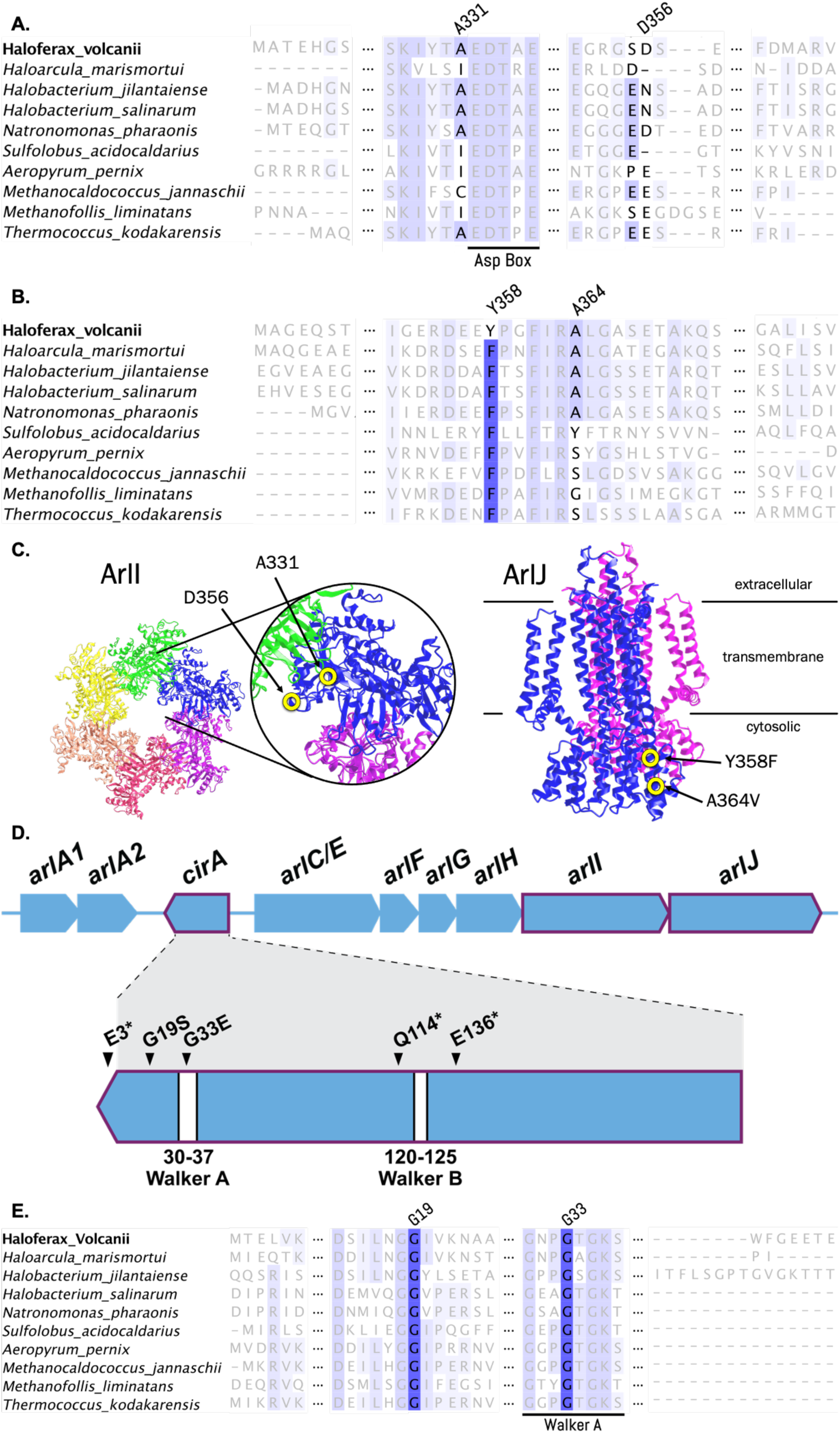
Residues of interest as sequence alignments, 3D structures, and genomic locations. Sequence alignments of **a)** ArlI, **b)** ArlJ and **e)** CirA from *Hfx. volcanii* with homologs from four additional haloarchaeal species and five non-halophilic archaeal species of different phyla. Regions of the alignment surrounding the mutated residues from **Table 1** are shown; other regions of the protein are either omitted or set to 25% transparency to ease visualization. Conserved residues across species are indicated in shades of blue, with darker blues corresponding to highly conserved residues. The ArlI conserved Aspartate Box motif and the CirA Walker A domain are indicated. **c)** Residues of ArlI and ArlJ corresponding to Δ*pilA*[*1-6*] suppressor mutations mapped to the AlphaFold structure predictions of the ArlI hexamer and ArlJ dimer, respectively. Yellow circles indicate the residues of interest. **d)** *cirA* is found between the archaellin genes (*arlA1*, *arlA2*) and the rest of the *arl* genes (*arlC/E–arlJ*). It is transcribed in the opposite direction as the *arl* genes. The genic locations of the residues corresponding to the mutations found in the Δ*pilA*[*1-6*] suppressor mutants as well as hypermotile mutants from a previous study^25^ are indicated, along with the conserved Walker A and Walker B motifs.

Mapping the two ArlI residues mutated in the suppressor mutants (**Table 1**), we found that the residue at position 331 of ArlI occurs just prior to the Aspartate Box, which is involved in divalent cation binding^29^. While ArlI A331 is not highly conserved across the species included in our alignment, the residues that align with *Hfx. volcanii* A331 are generally aliphatic (**Fig. 2a**). The replacement mutation ArlI^A331V^ was identified in two separately isolated suppressors, KY-1 and KY-8 (**Table 1**), suggesting the importance of alanine with regard to suppression of the Δ*pilA*[*1-6*] motility phenotype. *Hfx. volcanii* ArlI D356 is in a region with one or two negatively charged amino acids, including the glutamate at position 312 of *S. acidocaldarius* that is proposed to be part of a salt bridge between subunits of the ArlI hexamer^30^ (**Fig. 2a**).

Similarly, we mapped the single residue changes of ArlJ from the suppressor mutants in the protein sequence alignments for this membrane protein. The residue at position 358 in *Hfx. volcanii* ArlJ, which is mutagenized from a tyrosine to a phenylalanine residue in the suppressor strain KY-1 (**Table 1**), is a phenylalanine residue in all of the other species used in our alignment (**Fig. 2b**). The alanine at position 364 of ArlJ is conserved in the haloarchaeal sequences aligned. To determine the location of these mutations on the three-dimensional proteins, we generated AlphaFold structural predictions^31^ of the wild-type *Hfx. volcanii* ArlI hexamer and ArlJ dimer (**Fig. 2c**). The locations of the residues at position 331 and 356 on the ArlI structure suggest that they are potentially part of interaction sites with ArlJ or other interacting partners. Additionally, the *Hfx. volcanii* ArlI^D356^ residue overlaps with the corresponding residue E312 from the published structure of *S. acidocaldarius* FlaI (ArlI)^30^ (**Fig. S2a**) and may interact with *Hfx. volcanii* ArlI^H234^ to form a salt bridge (**Fig. S2b**). Thus, it is possible that the *Hfx. volcanii* ArlI^D356H^ mutation alters this interaction. For ArlJ, consistent with a potential interaction with cytoplasmic ArlI, the mutated residues appear not to affect the dimer interface and map to the cytoplasmic region as determined by SignalP 6.0^32^ (**Fig. 2c**).

Although *cirA* is found in the same genomic region as *arlI* and *arlJ*, it is transcribed in the reverse direction (**Fig. 2d**), and in contrast to the *arlI* and *arlJ* suppressors, the *cirA* suppressors harbor disruptive missense or nonsense mutations. One of these mutations results in a premature stop codon (CirA^W199*^) while the other two substitutions result in significant changes in the chemical properties (CirA^G19S^, CirA^G33E^) (**Table 1**). Protein sequence alignment of *Hfx. volcanii* CirA with the homologs of the same species used for ArlI and ArlJ revealed that G19 and G33 are highly conserved across species (**Fig. 2e**). The mutation for CirA^G33E^ appears within the conserved Walker A ATPase motif present in the KaiC-like domain (**Fig. 2d**). The premature stop codon and other disruptive mutations in *cirA* strongly support the idea that suppression of the Δ*pilA*[*1-6*] motility defect requires inactivation of CirA.

### The ArlI and ArlJ mutations suppress the **Δ***pilA*[*1-6*] motility defect and retain the archaella biogenesis function

To confirm that the mutated residues in ArlI and ArlJ contribute to the suppression of the Δ*pilA*[*1-6*] motility defect, we constructed Δ*arlI* and Δ*arlJ* strains as well as Δ*pilA*[*1-6*] strains also lacking either *arlI* or *arlJ*. The wild-type *arlI* and *arlJ* genes as well as *arlI* and *arlJ* genes with mutations corresponding to the suppressor mutants (**Table 1**) were inserted into the expression vector pTA963^33^. Each construct was transformed into the corresponding single deletion strain of Δ*arlI* or Δ*arlJ* as well as into the multi-deletion strains Δ*pilA*[*1-6*]Δ*arlI* or Δ*pilA*[*1-6*]Δ*arlJ*.

The Δ*arlI* and Δ*arlJ* strains are non-motile, but stab-inoculates of these strains overexpressing wild-type ArlI and ArlJ, respectively, on soft agar result in motility halos. The overexpression of ArlI^A331V^ or ArlI^D356H^ in the Δ*arlI* background also supports motility on soft agar, although the motility halo for the deletion strain expressing ArlI^D356H^ is smaller than that of ArlI or ArlI^A331V^ (**Fig. 3**). This supports the view that these mutations do not inactive the archaella biosynthesis function of ArlI. Similarly, the overexpression of either ArlJ^Y358F^ or ArlJ^A364V^ complements the Δ*arlJ* motility defect. In fact, overexpression of the point mutations results in slightly larger motility halos as compared to overexpression of ArlJ, indicating that these mutations also do not interfere with archaella biogenesis.

**Figure 3.**
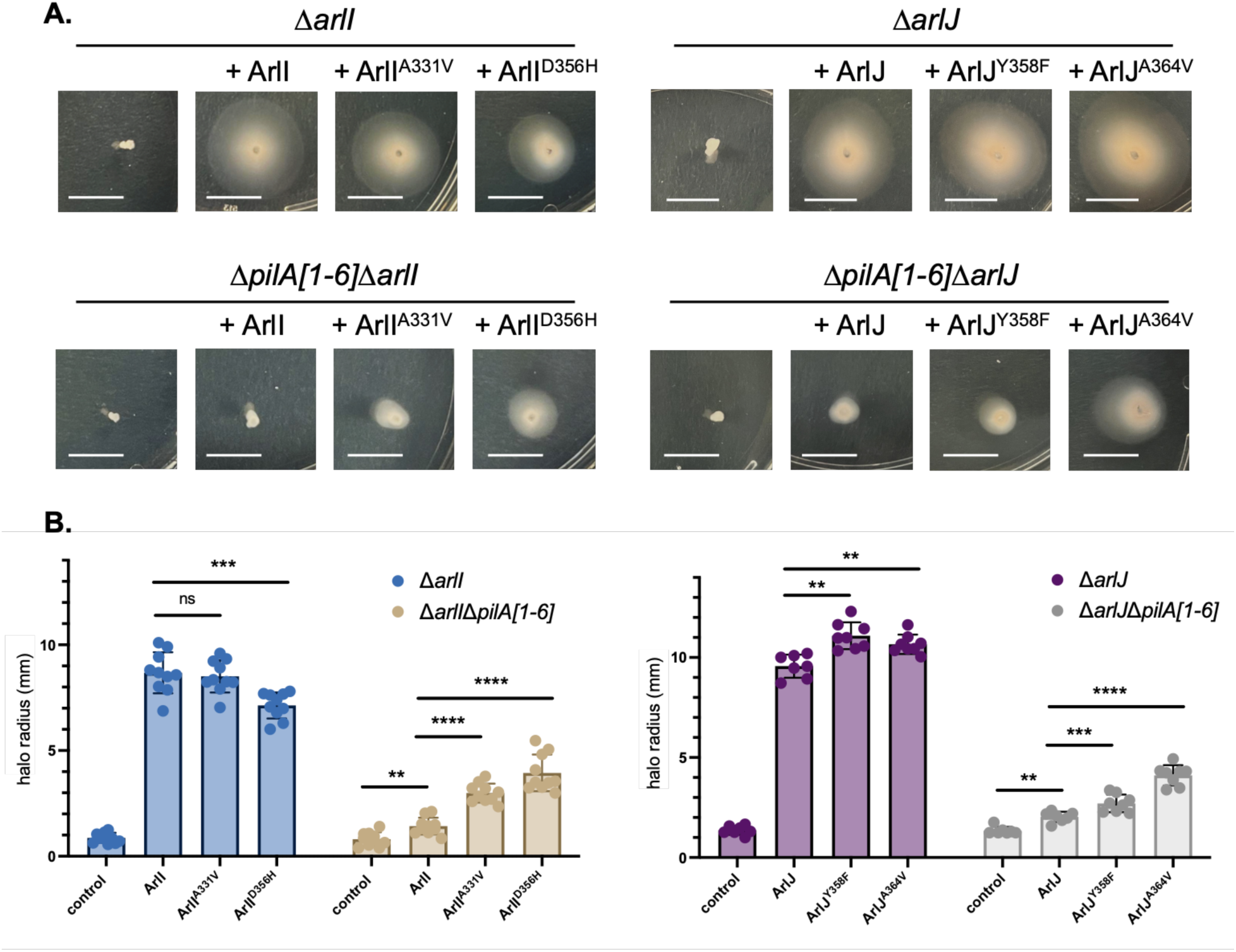
Overexpression of mutant ArlI or ArlJ results in differential motility. **a)** pTA963 expression vectors containing genes encoding either wild-type or mutant proteins were transformed into the indicated deletion strains. Empty vector was also transformed as control for all four deletion strains. Cells were stab-inoculated into 0.3% w/v agar plates, incubated at 45°C for 3 days and then at RT for 1 day before imaging. White bar represents 1 cm. **b)** Quantification of halo radius in millimeters across 7 to 10 replicates for each strain. Statistical analysis was performed using the unpaired nonparametric t-test. ** p<0.01, *** p<0.001 and **** p<0.0001.

Δ*pilA*[*1-6*]Δ*arlI* and Δ*pilA*[*1-6*]Δ*arlJ* are, as expected, non-motile. Overexpression of ArlI and ArlJ in the respective multi-deletion strains results in minimal motility, possibly due to the high abundance of plasmid-encoded ArlI or ArlJ, respectively. Supporting the importance of the point mutations, ArlI^A331V^ and ArlI^D356H^ overexpression results in significantly increased motility of the Δ*pilA*[*1-6*]Δ*arlI* strain. The Δ*pilA*[*1-6*]Δ*arlI* strain overexpressing ArlI^D356H^ has a larger motility halo than the strain overexpressing ArlI^A331V^, suggesting that the D356H mutation is more effective for overcoming the loss of pilins. Similarly, overexpression of ArlJ^Y358F^ and ArlJ^A364V^ in Δ*pilA*[*1-6*] Δ*arlJ* results in notable increases in the motility halo radius when compared to the overexpression of ArlJ in this strain. Additionally, ArlJ^A364V^ results in a larger motility halo than ArlJ^Y358F^, implying that it is more effective at suppressing the phenotype (**Fig. 3**). These results indicate that the single residue mutations in ArlI and ArlJ can enable motility in the absence of the conserved H-domain of the pilins.

### CirA suppresses swimming motility

Overexpression of ArlI or ArlJ in the multi-deletion strains led to a statistically significant increase in the motility halo radius compared to the negative control containing only the expression vector (**Fig. 3**), suggesting that increasing the abundance of ArlI or ArlJ can partially overcome pilin-mediated motility suppression. Since a) CirA is predicted to have regulatory roles^22^, b) *cirA* is found adjacent to the archaella genes (**Fig. 2d**), and c) three of the Δ*pilA*[*1-6*] motile suppressors have CirA disruptions along with missense mutations in ArlI and/or ArlJ (**Table 1**), we hypothesized that CirA regulates motility by affecting the abundance of ArlI and ArlJ.

To study the effects of disrupting CirA function, we generated the deletion strain Δ*cirA*. While Δ*cirA* is hypermotile, complementation with *cirA* expressed from pTA963 reverses this effect and significantly reduces the size of Δ*cirA* motility halos. Furthermore, the overexpression of CirA in wild-type *Hfx. volcanii* results in reduced motility (**Fig. 4**), suggesting that CirA plays a role in repressing motility.

**Figure 4.**
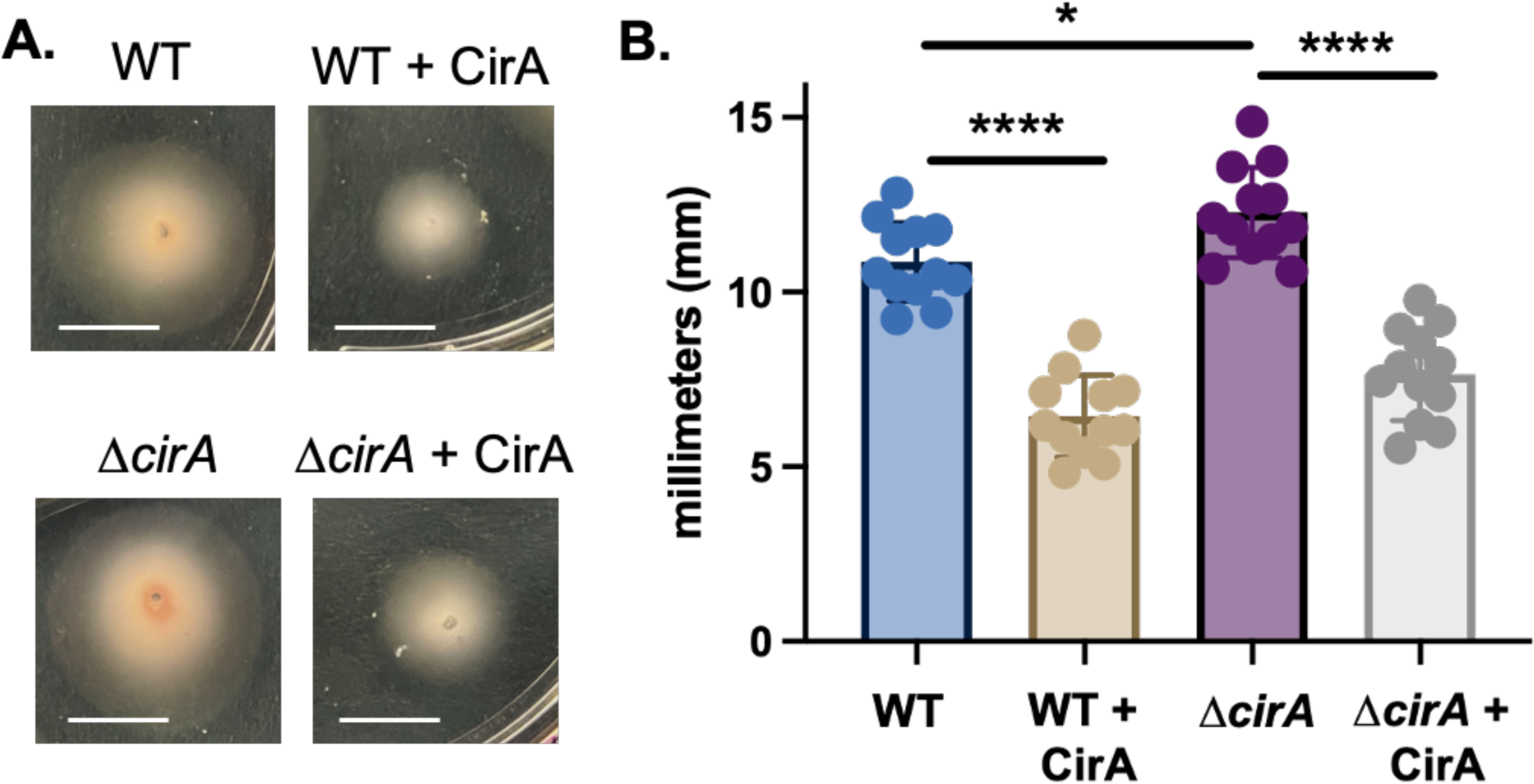
Overexpression of CirA results in reduced motility. **a)** Empty pTA963 or vector containing *cirA* was transformed into wild type and Δ*cirA* strains. Cells were stab-inoculated onto 0.3% w/v agar plates, incubated at 45°C for 3 days, then at RT for 1 day before imaging. White bar represents 1 cm. **b)** Quantification of halo radius across 12 replicates of each stabbed strain. Statistical analysis was performed using the unpaired nonparametric t-test. * p<0.05 and **** p<0.0001.

### Deletion of *cirA* results in smaller colonies, disk-deficiency, and decreased light scattering

Since KaiC-like ATPases are hypothesized to be major hubs of complex regulatory networks in archaea^22^, it follows that CirA could serve several regulatory roles in *Hfx. volcanii*. Consistent with this idea, the loss of CirA results in several phenotypic changes in this haloarchaeon. Δ*cirA,* compared to wild type, forms smaller, darker colonies when streaked on an Hv-Cab solid agar plate (**Fig. S3a**) and also produces a smaller, darker cell pellet upon centrifugation of mid-log phase cultures, grown to OD_600_ 0.55 (**Fig. S3b**). Using a plate reader equipped to monitor OD_600_ values, Δ*cirA,* after entering exponential phase, lagged by about three hours, equivalent to approximately one doubling time, compared to wild type (**Fig. S3c**). Additionally, since Δ*cirA* is hypermotile (**Fig. 4**), and motility is often associated with the rod shape^24^, we investigated Δ*cirA* cell shape. Interestingly, Δ*cirA* does not transition to disk-shaped cells at mid-log growth phase like wild type and remains as rod-shaped (**Fig. 5**). Considering the shape difference as well as the pellet size difference between Δ*cirA* and wild type, we wanted to determine whether the cell count of these cultures at the same OD_600_ is comparable. Surprisingly, colony forming units (CFU) counts of cultures grown to OD_600_ 0.5 yielded nearly four times the number of cells in Δ*cirA* compared to wild type (**Fig. S3d**) indicating that using optical density to measure *Hfx. volcanii* cell growth may not always accurately reflect doubling time.

**Figure 5.**
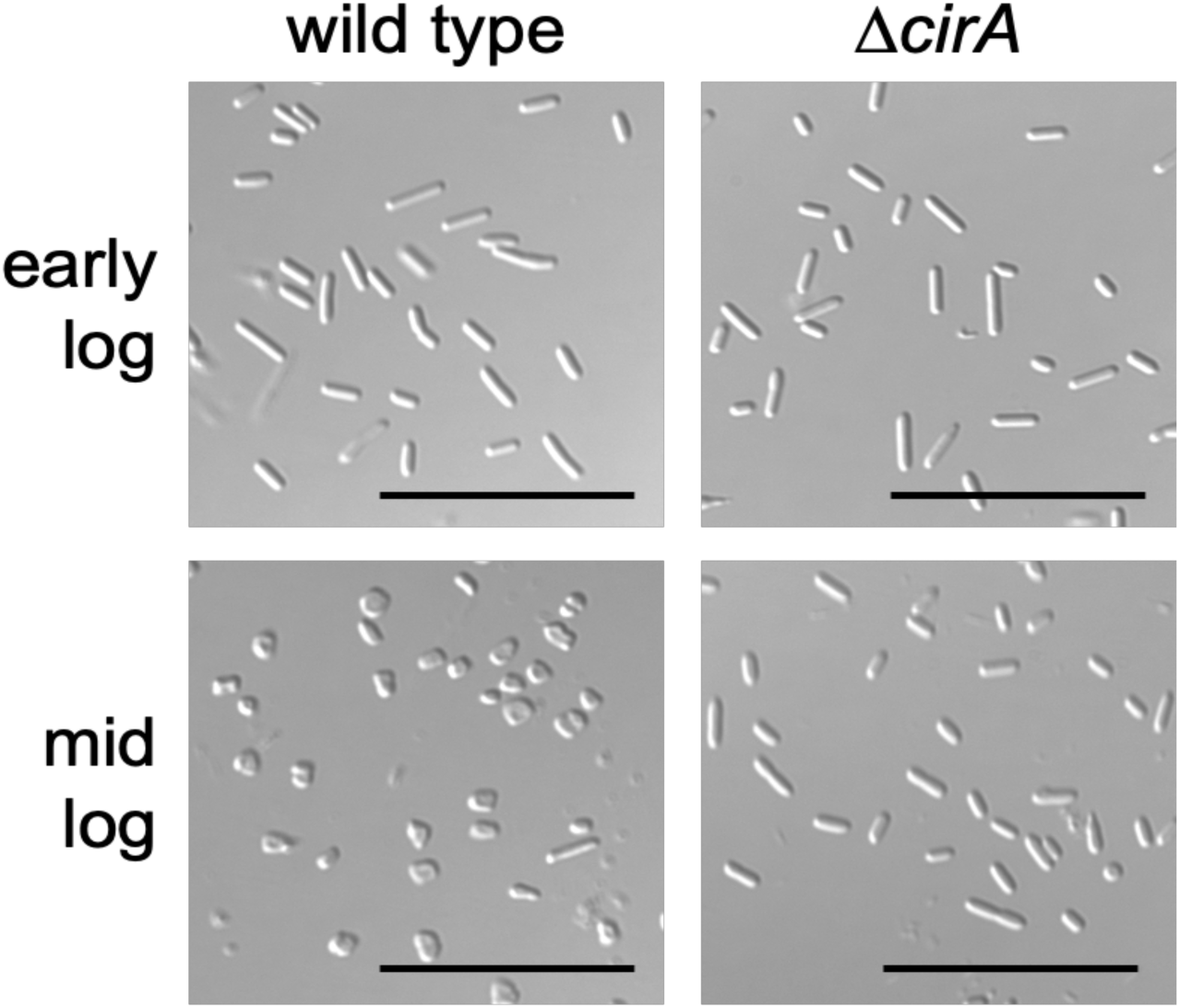
Δ*cirA* remains rod-shaped in mid-log phase. Early-log (OD600 0.05) and mid-log (OD600 0.35) phase cultures of wild-type H53 and Δ*cirA* were harvested and cells were imaged using differential interference microscopy (DIC). The black bar represents 20 µm.

### CirA regulates motility by reducing transcription of *arl* genes

Since overexpression of CirA results in reduced motility (**Fig. 4**), compounded with our observations that Δ*cirA* is hypermotile (**Fig. 4**) and has a prolonged rod stage (**Fig. 5**), we predicted that the loss of CirA affects the abundance of archaellins and the archaella biosynthetic components. Consistent with this hypothesis, the proteomics studies conducted by Schiller *et al.* determined that archaellin ArlA1 occurs at a higher abundance in early-log rod-shaped cells when compared to late-log disk-shaped cells^25^. While no differential abundance was observed for the biosynthetic components of the archaella, to date, proteomics has failed to identify ArlI or ArlJ peptides under any conditions tested^34^, suggesting low abundance of these biosynthetic components. Quantitative real-time polymerase chain reaction (qRT-PCR) studies, indeed, revealed that as wild-type cells begin the transition from rod-shaped in early-log to disk-shaped in mid-log, the transcription of genes encoding the archaellin (ArlA1) and archaella biosynthetic components (ArlI and ArlJ) decreases (**Fig. 6**). Transcription levels of all three genes are higher in Δ*cirA* early-log when compared to wild type early-log, and the higher levels are maintained as Δ*cirA* transitions into mid-log. Thus, *cirA* plays a role in the transcription of the *arl* genes.

**Figure 6.**
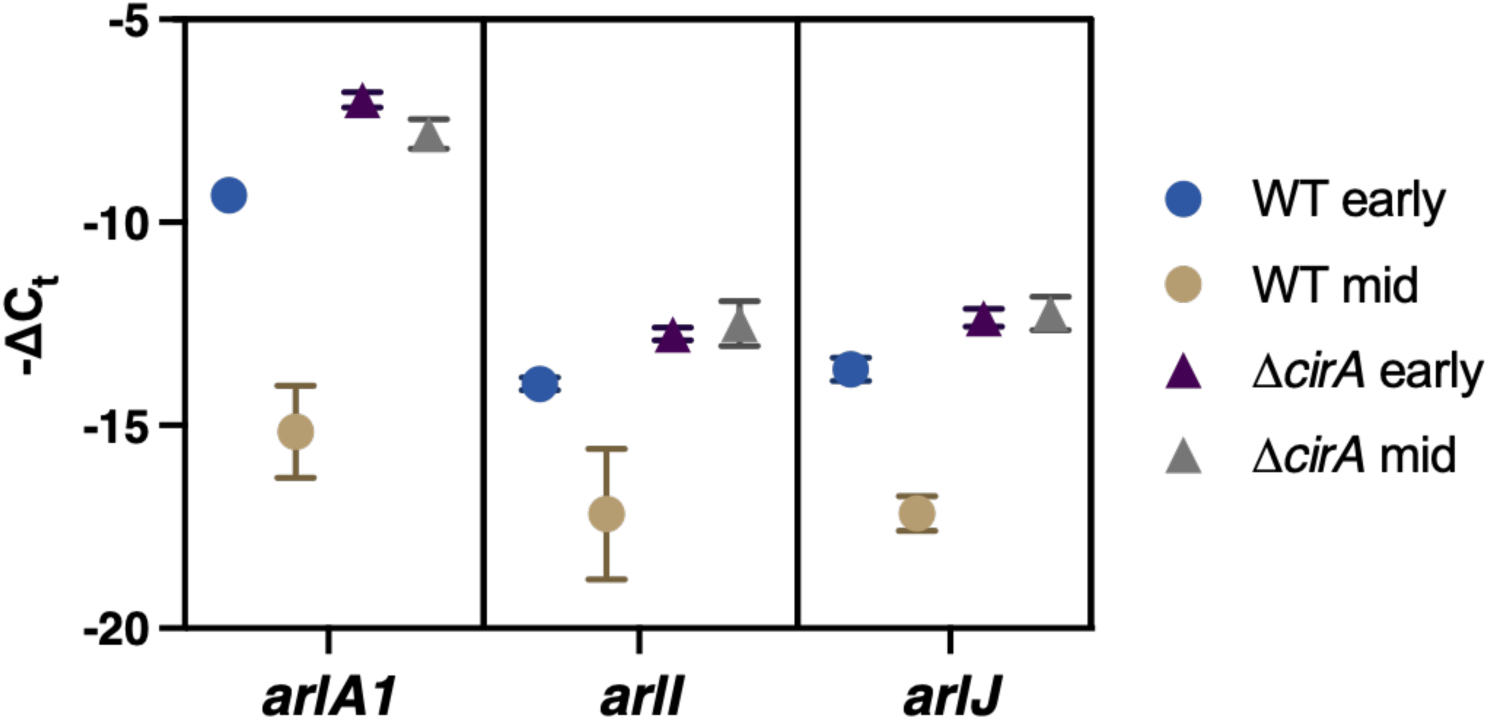
Δ*cirA* displays a different transcription profile of *arlA1*, *arlI*, and *arlJ* than wild type. qRT-PCR was conducted using primers for 16S rRNA, *arlA1*, *arlI*, and *arlJ* under the following conditions: 1) wild type at early-log; 2) wild type at mid-log; 3) Δ*cirA* at early-log; and 4) Δ*cirA* at mid-log. Values are shown as the negative difference in threshold crossing point –ΔCt of *arlA1*, *arlI*, and *arlJ* relative to 16S rRNA. Error bars represent the standard deviation of five technical replicates.

## Discussion

It has been proposed that during biofilm formation, *Hfx. volcanii* rapidly assembles membrane-sequestered pilins into type IV pili in order to promote surface adhesion. Thus, the pilins are depleted from the membrane, which leads to the cessation of motility^27^. This loss of motility due to a depletion of membrane-sequestered pilins is presumably mimicked by the Δ*pilA*[*1-6*] strain. In this study of the mutations that arose in motile suppressors of the Δ*pilA*[*1-6*] strain, we have discovered a novel aspect of *Hfx. volcanii* motility regulation. The identification of several suppressor mutations in genes encoding ArlI and ArlJ suggested that by altering specific residues in these archaella biosynthetic components, motility can be restored to the non-motile Δ*pilA*[*1-6*] strain. Indeed, we revealed with reverse genetics that the mutated proteins, which can complement the respective *arlI* and *arlJ* deletion strains, can restore motility to the multi-deletion strains Δ*pilA*[*1-6*]Δ*arlI* and Δ*pilA*[*1-6*]Δ*arlJ* (**Fig. 3a**). Most of these mutations are point mutations in *arlI* and *arlJ* that result in single residue changes which retain the same chemical properties as the replaced residue. Therefore, it is possible that these slight variations of key residues alter binding affinity(s) of these proteins to other proteins. Alignments of ArlI reveal that the D356 residue aligns with a region that contains one or two negatively charged amino acids in the other homologs, including an aspartate in *S. acidocaldarius* that forms a putative salt bridge between subunits of the hexamer^30^. In *Hfx. volcanii* ArlI, D356 may form a salt bridge with H234 (**Fig. S2b**), an interaction which seems to be disrupted in ArlI^D356H^. The disruption of this putative salt bridge may explain the reduced motility of the overexpression of ArlI^D356H^ compared to that of ArlI and ArlI^A331V^ in the Δ*arlI* strain (**Fig. 3**). The increased motility of the overexpression of ArlI^D356H^ in Δ*pilA*[*1-6*]Δ*arlI* may still be due to the mutation altering binding affinity to partner proteins. Mutations in ArlJ do not suggest their involvement in dimer production but their cytoplasmic location may indicate interactions with cytoplasmic proteins, such as ArlI, suggesting that binding affinity for a partner protein at the cytoplasmic interface is altered. In the case that the point mutations in ArlI and ArlJ lead to increased binding affinity, the interactions between binding partners that are necessary for motility could be increased, and thus the mutants are able to synthesize archaella despite missing the conserved pilin H-domain. Conversely, if the point mutations lead to decreased binding, it is possible that the mutant proteins no longer bind to inhibitory proteins that suppress motility and become capable of swimming without the H-domain. Further analysis of the interactome of ArlI and ArlJ through protein-protein interaction studies may elucidate additional components involved in pilin-mediated motility regulation.

In the motile suppressors of Δ*pilA*[*1-6*], mutations in *cirA* arose concurrently with several mutations in the archaella biosynthetic components (**Table 1**). A potential link between CirA and hypermotility has been alluded to in a previous study, where an *Hfx. volcanii* transposon mutant (JK3) had been identified as hypermotile. While the gene disrupted by the transposon did not lead to hypermotility, WGS of the transposon mutants uncovered a secondary mutation in *cirA*^25^. The secondary mutation resulted in a premature stop codon at the third amino acid (CirA^E3*^). Similarly, in the Δ*pilA*[*1-6*] suppressors, the *cirA* mutation in KY-7 results in a premature stop codon, and the KY-1 and KY-8 strains harbor disruptive *cirA* point mutations in high conserved residues (**Fig. 2e**). Here we show that a Δ*cirA* strain is indeed hypermotile, potentially due to the lack of CirA-dependent *arl* gene regulation.

In our previous study, we discovered, using quantitative proteomics, that the archaellin ArlA1 has a significantly lower abundance in late-log phase than early-log phase, while CirA increases in abundance across the growth phases^25^. Interestingly, ArlA1 abundance is not as drastically different in the growth phase comparisons of JK3, suggesting that CirA is involved in the regulation of ArlA1. While differences in protein abundance of ArlI or ArlJ were not observed in wild type or JK3^25^, the abundance of archaella biosynthetic components could be below the detection level of quantitative proteomics experiment, consistent with the absence of any ArlI or ArlJ peptides identified in the Archaeal Proteome Project (ArcPP) database^34^. However, a study in another halophilic species found that while protein abundance changes for the archaella biosynthetic proteins in the tested experimental conditions were not found, transcriptional analysis was able to identify differences in the regulation of *arl* genes^35^. Furthermore, another study reported that sessile, disk-shaped biofilm-derived cells can exhibit a decreased expression of *arl* genes^36^. Corroborating those findings, in this study, we detected decreases in the transcription levels of not only *arlA1*, but also of *arlI* and *arlJ*, between early-log phase rods and mid-log phase disks in wild type. This decrease was not observed in the Δ*cirA* strain (**Fig. 6**), supporting our hypothesis that CirA is involved in the downregulation of *arlA1*, *arlI*, and *arlJ*. It is yet unknown whether CirA functions directly as a regulator of the transcription or indirectly as an intermediary in a regulatory network. CirA is transcribed in the opposite direction as the *arl* genes (**Fig. 2d**), but it is unlikely that the act of transcribing CirA reduces the transcription of the *arl* genes because the overexpression of CirA from a plasmid also reduces motility (**Fig. 4**). It is also important to note that the loss of CirA affects more than just motility, including the inability to form disks in the conditions tested (**Fig. 5**) and the formation of smaller colonies (**Fig. S3a**). Additionally, pellets of Δ*cirA* cells in liquid cultures grown to the same OD_600_ as wild type are also smaller even though viability counts reveal strikingly higher cell numbers (**Fig. S3b,d**), suggesting that CirA affects the overall density of the cells, potentially by modulating cell surface composition. Interestingly, the OD_600_ growth curve did reveal that both wild type and Δ*cirA* cells reach a similar optical density at stationary phase (**Fig. S3c**). While more in-depth studies of cell culture growth of wild type and Δ*cirA* are needed, the multiple physiological differences between these two strains observed in this study supports that CirA is a broad regulatory protein in *Hfx. volcanii*. Expression of *cirA* has been shown to oscillate in a light/dark dependent manner, reminiscent of the KaiC regulator. In fact, one of the suppressor mutations, CirA^G33E^, occurs within the Walker A motif, a region which has previously been shown to induce an arrhythmic phenotype when mutated^37^. Further characterization of CirA function will elucidate the mechanism by which it suppresses motility, affects physiology, and potentially modulates *Hfx. volcanii* light-dependent behavior.

Lastly, additional mutations in the Δ*pilA*[*1-6*] suppressor mutants may also play a role in motility regulation. In particular, KY-2 and KY-6 lack mutations in *arlI*, *arlJ*, or *cirA* (**Table S1**). Repeating our motility screen could help narrow down additional important components to further help elucidate mechanisms of motility regulation in *Hfx. volcanii*.

In conclusion, in this study we used the model organism *Hfx. volcanii* to uncover that ArlI and ArlJ are involved in pilin-mediated motility regulation and identified critical residues important to this regulatory function. Furthermore, we have determined a novel role for CirA in motility regulation and revealed its importance in disk formation and cell density.

## Materials and Methods

### Growth conditions

The plasmids and strains used in this study are listed in **Table S2 and S3**. *Hfx. volcanii* H53, H98, and their derivatives were grown at 45°C in liquid (orbital shaker at 250 rpm, 1-in orbital diameter) or on solid agar (1.5% w/v) in either semi-defined Casamino Acid (Fisher Scientific) Hv-Cab medium^23^ supplemented with uracil (50 µg mL^-1^ final concentration, Sigma) or Modified Growth Medium (MGM)^38^. Hv-Cab requires further supplementation with tryptophan (50 µg mL^-1^ final, Fisher Scientific) or with thymidine (40 µg mL^-1^ final, Acros Organics) and hypoxanthine (40 µg mL^-1^ final, Sigma) for H53 and derivatives or for H98 and derivatives, respectively. MGM requires no additional supplementation. Upon transformation with plasmid constructs that encode *pyrE2* or *hdrB*, supplementation with uracil or thymidine/hypoxanthine in Hv-Cab is no longer required. Solid agar plates are placed within sealable bags during incubation to prevent the plates from drying out. *Escherichia coli* strains were grown at 37°C in NZCYM (RPI) medium supplemented with ampicillin (100 µg mL^-1^ final, Corning)^39^.

### Ethyl methanesulfonate (EMS) mutagenesis and motility screen

The Δ*pilA*[*1-6*] strain was inoculated in MGM and grown to stationary phase (OD_600_ 1.0). The cultures were then diluted with fresh MGM at 1:10. The sub-inoculation was then grown until OD 0.3-0.4. Two mL aliquots of the culture were transferred to five microcentrifuge tubes corresponding to five experimental conditions: the negative control with no EMS exposure as well as 0, 10, 20, and 30 minutes of EMS exposure. Cells were centrifuged and washed twice with SMT buffer (3.5 M NaCl, 0.15 M MgSO4, 10 mM Tris-Cl, pH 7.2) and then resuspended with SMT buffer. 10.29 µl EMS (0.05 M) was added to the tubes, mixed thoroughly, and incubated for the corresponding length of exposure time. Cells were then centrifuged and washed twice with SMT buffer. Using a toothpick, cell pellet was touched and stab lines were made into 0.35% agar MGM plates. Plates were incubated upright at 45°C for 3-5 days until small halos extending from the stab lines started to appear. Motility phenotype was confirmed by using a toothpick to touch the halo, re-streak onto solid agar (1.5% w/v), and re-stab single isolated colonies onto 0.35% w/v agar MGM plates.

### Sequencing **Δ***pilA*[*1-6*] suppressor strains

The methods for cell cultivation, DNA extraction, and sequencing were the same as described previously for mutants derived from the H295 strain^40^, but here the parent strain was the H53 strain of *Hfx. volcanii* (Δ*pyrE2*, Δ*trpA*), which similarly has the pHV4 plasmid integrated into the chromosome and lacks pHV2. The sequence reads (Illumina, paired-end, 150 cycle reads) for the suppressor strains were imported into Geneious Prime (v. 2023.2.1), trimmed using BBDuk (v. 38.84; default settings) from the BBTools package^41^, and mapped to the genome sequence of the parental strain (H53) using the Bowtie2 mapper (v. 2.4.5). The mapped assemblies were scanned manually within Geneious in order to detect all differences across the entire genome of each strain. Mutations for each strain are listed in **Table S1**.

### Protein amino acid sequence alignments

Protein sequences of *Hfx. volcanii* DS2 ArlI, ArlJ, and CirA were found on the UniProt protein database and a protein BLAST search was performed of microbial genomes within GenBank on the NCBI website. Four haloarchaea species (*Haloarcula marismortui, Halobacterium jilantaiense, Halobacterium salinarium,* and *Natronomonas pharaonis*) and five other representative archaeal species from different phyla (*Sulfolobus acidocaldarius, Aeropyrum pernix, Methanocaldococcus jannaschii, Methanofollis liminatans,* and *Thermococcus kodakarensis*) were chosen for the sequence alignment. Sequences were aligned via Clustal Omega Multiple Sequence Alignment^42^ and the output URL was imported to Jalview (v. 2.11.3.2)^43^. Aligned residues were colored on Jalview according to “Percentage Identity” and the program automatically assigned darker blue colors residues to indicate a higher percentage of conservation. Residues corresponding to the point mutations of *Hfx. volcanii* ArlI, ArlJ, and CirA from **Table 1** were labeled. The conserved motifs for ArlI proteins which had previously been identified in *S. acidocaldarius* ArlI (Walker A and Walker B nucleotide binding ATPase sites, the aspartate box, the histidine box)^29^ were identified and indicated in **Fig. S1**. The conserved motifs for *Hfx. volcanii* CirA (Walker A and Walker B domains, catalytic glutamate residues)^26^ were identified and indicated in **Fig. S1**.

### Protein structure prediction and modeling

The hexameric structure of the ArlI complex was predicted using a local installation of AlphaFold-Multimer^31^. AlphaFold-Multimer simulations were performed on the Wistar Institute High Performance Cluster (HPC). Five structures were predicted without a template, and the top ranked structure in terms of ipTM+pTM score was inspected using Pymol 2.5.5. The structure of the ArlJ complex was generated using AlphaFold (v. 2.3.1)^44^ via LocalColabFold (v. 1.5)^45^ using the default settings. Computations were performed on the Penn Arts and Sciences General Purpose Cluster (GPC). Structure files from both ArlI and ArlJ were imported into the iCN3D web-based 3D-structure viewer^46^ for visualization, color adjustments, and locating the residues of interest from **Table 1**. Transmembrane regions of the ArlJ dimer were determined by SignalP 6.0^32^ and the orientation of the protein complex was adjusted with respect to the extracellular and cytoplasmic regions.

### Plasmid preparation and *Hfx. volcanii* transformation

The pTA131 plasmid was used to generate chromosomal deletions^47^ and the pTA963 plasmid was used for complementation and overexpression experiments^33^. DNA Phusion-Taq polymerase, restriction enzymes, and DNA ligase were purchased from New England BioLabs. Plasmids were initially transformed into *E. coli* DH5α cells; plasmids were then transformed into *E. coli* DAM^−^ strain DL739 prior to *Hfx. volcanii* transformation. Plasmid preparations for both *E. coli* strains were performed using the PureLink™ Quick Plasmid Miniprep Kit (Invitrogen).

*Hfx. volcanii* transformations were performed using the polyethylene glycol (PEG) method^38^. All oligonucleotides used to construct the recombinant plasmids are listed in **Table S4**.

### Generation of chromosomal deletions

Chromosomal deletions were generated by the homologous recombination method (pop-in/pop-out), as previously described^47^. Plasmid constructs for use in the pop-in/pop-out deletion process were generated by using overlap PCR, as previously described^48^: about 750 nucleotides flanking the corresponding gene were amplified via PCR (for primers **Table S4**). This resulted in an overlapping region between the upstream and downstream fragments, which were subsequently fused by overlap PCR. The fused upstream and downstream fragments were then cloned into the haloarchaeal suicide vector pTA131 via digestion with XbaI and XhoI followed by ligation^47^. For the *cirA* deletion construct, XbaI and HindIII were used because *cirA* has an internal XhoI site. The resulting constructs were verified by Sanger sequencing. The knockout plasmid constructs for Δ*arlI,* Δ*arlJ,* and Δ*cirA* were transformed into H53 wild type to generate the single deletion strains. Additionally, the plasmid constructs for Δ*arlI* and Δ*arlJ* were transformed into the Δ*pilA[1-6]* background^27^ to generate multi-deletion strains. Pop-out transformants were selected on agar plates containing 5-fluoroorotic acid (FOA) (Toronto Research Chemicals Inc.) at final concentration 50 µg mL^-1^. Successful gene deletion was confirmed by PCR using primers within the deleted gene as well as the upstream and downstream primers used for initial plasmid construction. Deletion strains were further analyzed by Illumina whole genome sequencing and variant calling performed by SeqCenter (Pittsburgh, PA, USA) to confirm the complete deletion of the gene as well as search for any secondary genome alteration. Ilumina sequencing libraries were prepared using the tagmentation-based Illumina DNA Prep kit and custom IDT 10bp unique dual indices (UDI) with a target insert size of 280 bp. No additional DNA fragmentation or size selection steps were performed. Illumina sequencing was performed on an Illumina NovaSeq X Plus sequencer, producing 2×151bp paired-end reads. Demultiplexing, quality control and adapter trimming was performed with bcl-convert1 (v4.2.4). Variant calling was carried out using BreSeq (v0.38.1) under default settings using *Hfx. volcanii* DS2 (GCF_000025685.1) as a reference.

### Construction of expression plasmids

For gene complementation studies, overexpression of *arlI*, *arlJ*, or *cirA* were performed using pTA963, which includes a tryptophan-inducible promoter (p.*tna*) for the inserted genes^33^. The primers to amplify out the respective gene are listed in **Table S4**. The reverse primers were designed to incorporate a 6xHis-Tag to the end of the gene to enable expression of proteins for downstream protein purification applications. For *arlI* and *arlJ*, the PCR-amplified genes were ligated into pTA963; both the vector and the amplified gene fragment were digested with BamHI and EcoRI prior to ligation. The resulting constructs were verified by Sanger sequencing. For complementation, deletion strains were transformed with the plasmid containing the corresponding gene or with empty vector pTA963 as a control.

### Motility plate assay

Motility assays were assessed on 0.35% w/v agar Hv-Cab medium with supplements as required. A toothpick was used to stab-inoculate the agar followed by incubation at 45°C upright in a plastic box with added wet paper towels to maintain humidity. Paper towels were changed daily. Motility assay plates were removed from the 45°C incubator after three days and imaged after one day of room temperature incubation as this increased the carotenoid production, allowing for better visibility of the halos. Motility halos were quantified using Fiji (ImageJ) (v. 2.9.0)^49^. Images were uploaded to Fiji, and the scale was set based on plate diameter. Statistical significance of halo diameters was assessed with an unpaired t-test using GraphPad Prism v.9.5.1 (528) for macOS (GraphPad Software, San Diego, California USA, www.graphpad.com).

### Cell pellets

Liquid cultures in Hv-Cab medium with supplements as required of *Hfx. volcanii* wild type and Δ*cirA* were grown until OD_600_ 0.55. One ml of each culture was centrifuged in 1.5 ml Eppendorf tubes at 4,900 x g for six minutes and supernatant was aspirated out. Resulting pellets were imaged.

### Growth curve

Growth curves for H53 wild type and Δ*cirA* strains were measured using BioTek Epoch 2 microplate reader with BioTek Gen6 v.1.03.01 (Agilent). Colonies of each strain were inoculated in 5 mL Hv-Cab liquid medium and grown shaking at 45°C until an OD_600_ 0.15 was reached. Cultures were diluted to an OD_600_ 0.01 with fresh liquid medium in a final volume of 200 µL in each well of a non-treated and flat-bottom polystyrene 96-well plate (Corning). Eight biological replicates were used for each strain. To help prevent evaporation, two rows of perimeter wells were aliquoted with 200 µL of media and a two-degree gradient was specified in the 45°C temperature setting. Readings were taken every 30 minutes with double orbital, continuous fast shaking (335 cpm, 4 mm) in between. Readings were taken at a wavelength of 600 nm for a total of 48 hours and then plotted in GraphPad Prism v. 9.5.1 (528) for macOS (GraphPad Software, San Diego, California USA, www.graphpad.com).

### Colony forming unit (CFU) counting

5 mL liquid cultures of the H53 wild type and Δ*cirA* strains were inoculated in liquid Hv-Cab medium and grown to OD_600_ 0.5. 30 µL of a 10^-5^ dilution was spread onto Hv-Cab solid agar plates in three technical replicates and incubated for 5 days. Images were taken and CFUs were counted using the Fiji (ImageJ) (v. 2.9.0) “Cell Counter” feature^49^. Statistical significance was analyzed with an unpaired t-test using GraphPad Prism v. 9.5.1 (528) for macOS (GraphPad Software, San Diego, California USA, www.graphpad.com).

### Cell shape imaging

5 mL liquid cultures of the H53 wild type and Δ*cirA* strains were inoculated and incubated, with shaking, at 45°C until OD_600_ 0.05 and 0.35 were reached, corresponding to early and mid-log growth phase, respectively. 1 mL from each culture was centrifuged at 4,900 x g for six minutes. Supernatant was aspirated to leave ∼5 µL liquid remaining with the pellet for the cultures at OD_600_ 0.05 (200 times concentrated) and ∼100 µL liquid remaining with the pellet for the cultures at OD_600_ 0.35 (10 times concentrated). Pellets were resuspended in the corresponding remaining liquid and 1.5 µL of resuspension was placed on glass slides (Fisher Scientific) and a glass coverslip (Globe Scientific Inc.) was placed on top. Slides were visualized using a Leica Dmi8 inverted microscope attached to a Leica DFC9000 GT camera with Leica Application Suite X (v. 3.6.0.20104) software. Differential interference contrast (DIC) images were captured at 100x magnification.

### Quantitative RT-PCR

5 mL liquid cultures of H53 wild type and Δ*cirA* strains were inoculated and incubated, shaking, at 45°C until OD_600_ 0.05 and 0.35 were reached, corresponding to early and mid-log growth phase, respectively. 4 mL of the early-log cultures and 0.5 mL of the mid-log cultures were harvested, and pelleted at 4,900 x g for six minutes. RNA was extracted using the Qiagen RNeasy Plus Micro Kit and lysate homogenized with Qiagen QIAshredder columns. RNA concentration determined with Qubit™ RNA High Sensitivity (HS) Assay Kits. 2 µg of RNA from each condition was treated with 2 units of Promega RQ1 RNase-free DNase, incubated at 37°C for 45 minutes, and then DNase was heat inactivated. cDNA was synthesized from RNA using ProtoScript II First Strand cDNA Synthesis Kit (New England BioLabs) using random primers (included with kit). PrimeTime™ qPCR Primers for *Hfx. volcanii arlA1*, *arlI*, *arlJ*, and 16S rRNA were designed using Integrated DNA Technologies (IDT) (**Table S4**). qRT-PCR reaction mixtures were set up with 10 µl of SYBR® Green JumpStart™ *Taq* ReadyMix™ (Sigma-Aldrich), 2 µl primers (final concentration 1 µM), 1 µl of cDNA from each experimental condition, and nuclease-free water for a total reaction volume of 20 µl. Each reaction was performed with 5 technical replicates on QuantStudioLJ3 Real-Time PCR System using the ΔΔC_t_ experiment type. The thermal cycling conditions were as follows: initial denaturation 94°C for 2 min, then 40 cycles of 94°C for 15 s and 60°C for 1 min. The SYBR Green fluorescence signal was recorded at the end of each cycle and the cycle threshold (C_t_) was automatically determined by the instrument. Differential expression was calculated from the negative difference in threshold crossing point (–ΔC_t_) between the primers of interest (targeting *arlA1*, *arlI*, *arlJ*) and control primers that target 16S rRNA. –ΔC_t_ values were plotted in GraphPad Prism v. 9.5.1 (528) for macOS (GraphPad Software, San Diego, California USA, www.graphpad.com).

## Data Availability

Raw sequencing results for WGS of the Δ*pilA*[*1-6*] suppressor strains and raw values for the growth curve and qRT-PCR can be found at 10.5281/zenodo.10735612. All other data types associated with this manuscript can be found in the supplemental material.

## Acknowledgements

We gratefully acknowledge Wil Prall for his assistance with the qRT-PCR data analysis. We also thank Dr. Alex Bisson, Dr. Friedhelm Pfeiffer, and Yirui Hong for their insightful comments on this manuscript. This work was supported by the National Science Foundation Grant NSF- MBC2222076 and the University of Pennsylvania Research Fund grant. Priyanka Chatterjee was additionally supported by a National Institutes of Health Training Grant T32 GM007229 “Cell and Molecular Biology”. Marco Garcia was supported by a Merck/PennFERBS fellowship.

