## Supplemental Figures for "Involvement of ArlI, ArlJ, and CirA in Archaeal Type-IV Pilin-Mediated Motility Regulation"

### Supplemental Figures and Tables

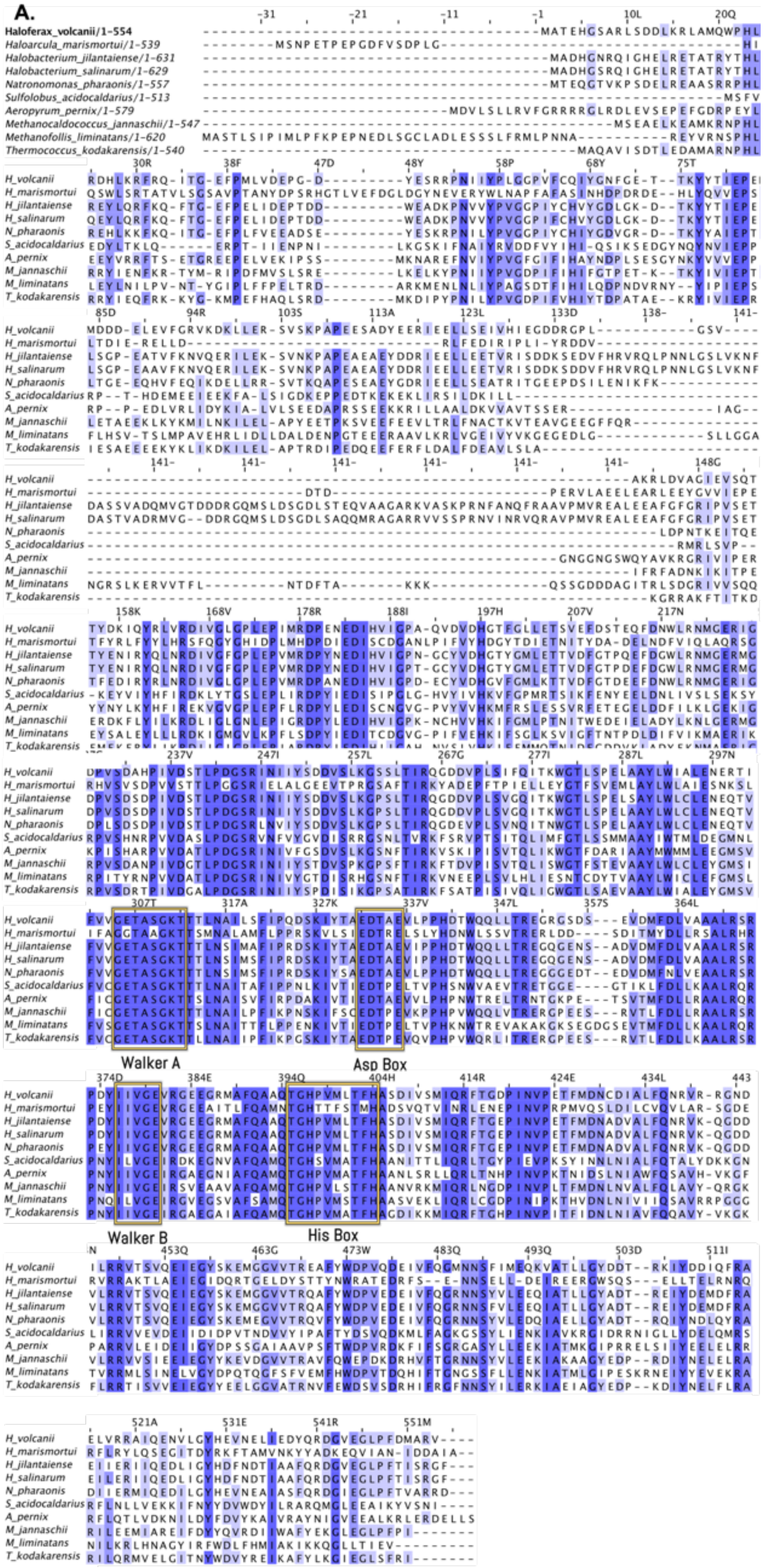

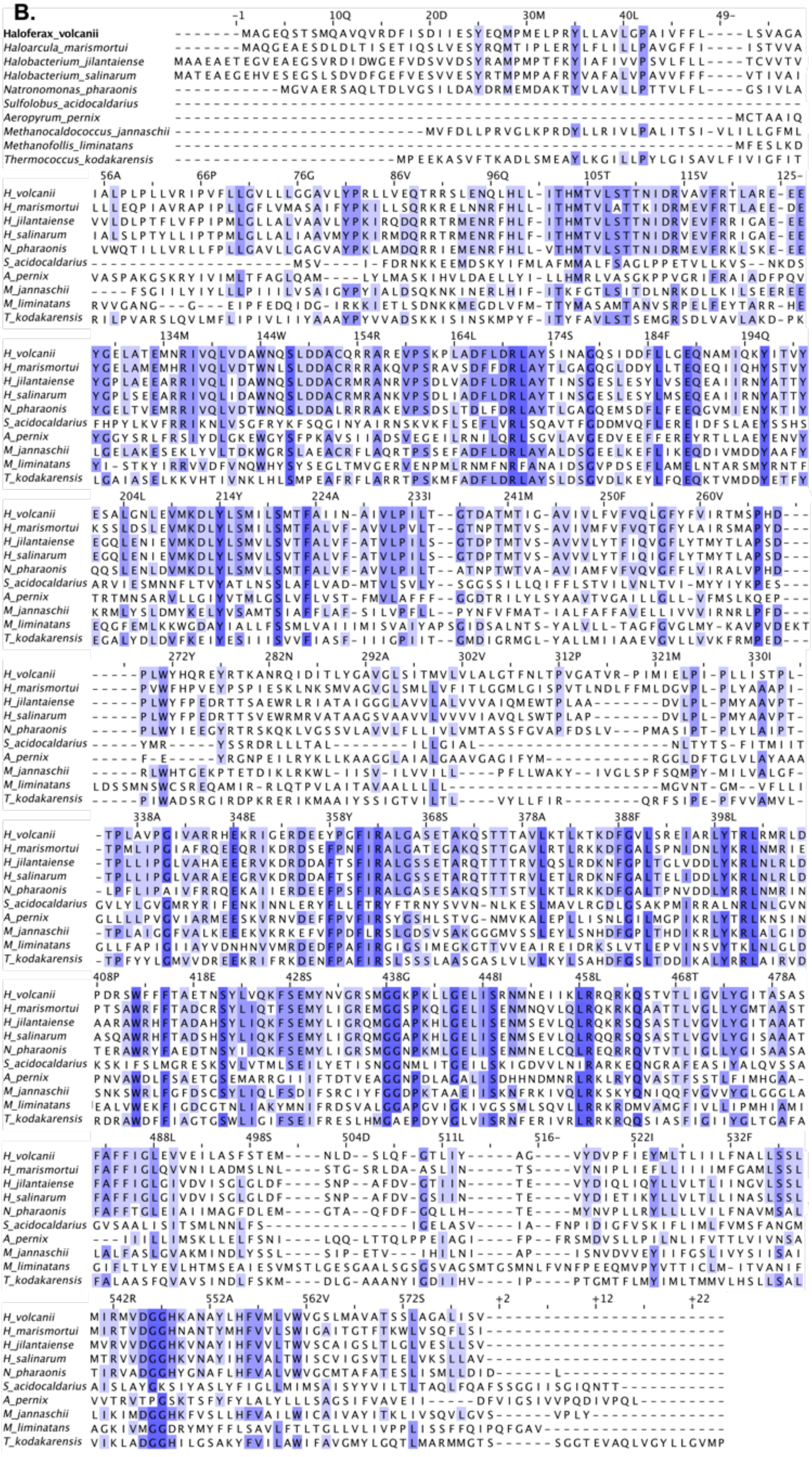
B.
Haloferax_volcanii
Haloarcula_marismortui
Halobacterium_jilantaiense
Halobacterium_salinarum
Natronomonas_pharaonis
Sulfolobus_acidocaldarius
Aeropyrum_pernix
Methanocaldococcus_jannaschii
Methanofollis_liminatans
Thermococcus_kodakarensis

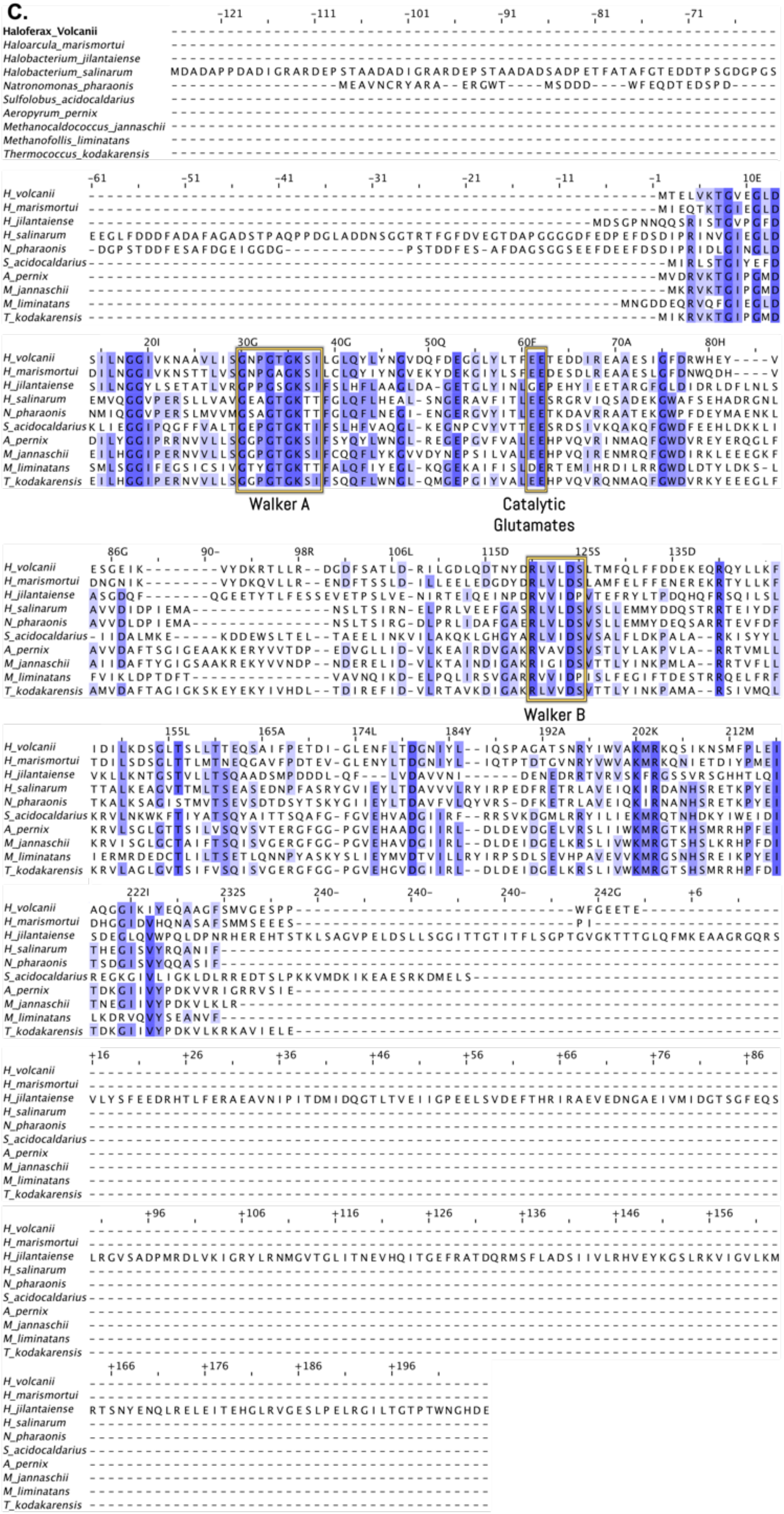
C.
Haloferax_Volcanii
Haloarcula_marismortui
Halobacterium_jilantaiense
Halobacterium_salinarum
Natronomonas_pharaonis
Sulfolobus_acidocaldarius
Aeropyrum_pernix
Methanocaldococcus_jannaschii
Methanofollis_liminatans
Thermococcus_kodakarensis
Walker A
Catalytic Glutamates
Walker B

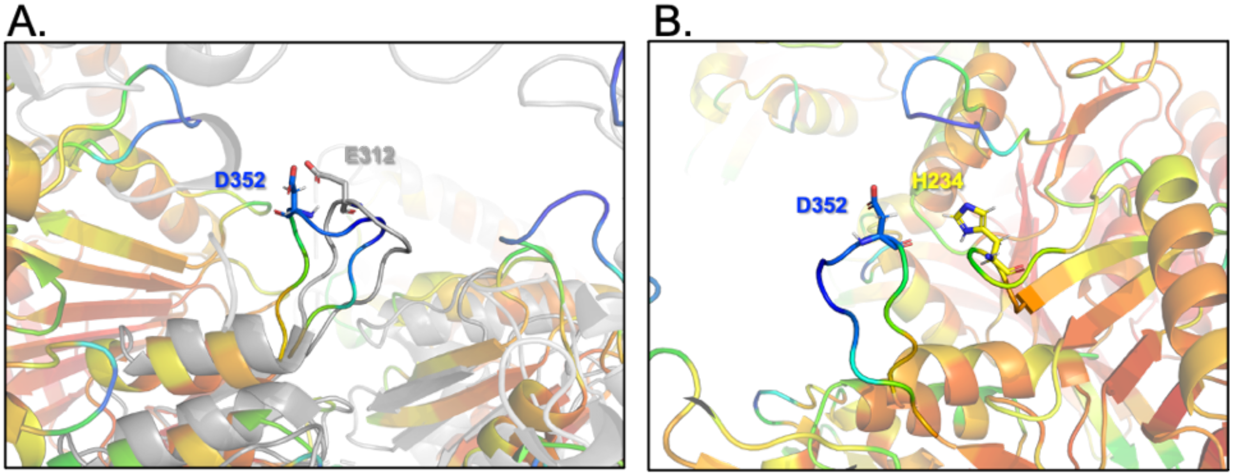


**Figures S2**

**Potential interactions of residue D352. a)** An alignment of the crystal structure of the published structure of *S. acidocaldarius* ATPase FlaI (ArlI)[30] (PDB: 4II7, grey) to the predicted *Hfx. volcanii* ArlI hexamer model (colored red to blue by decreasing pLDDT). Homologous positions between the *Hfx. volcanii* residue D352 and *S. acidocaldarius* residue E312 are shown. **b)** *Hfx. volcanii* ArlI residue D352 is proximal to residue H234 on the neighboring protomer.

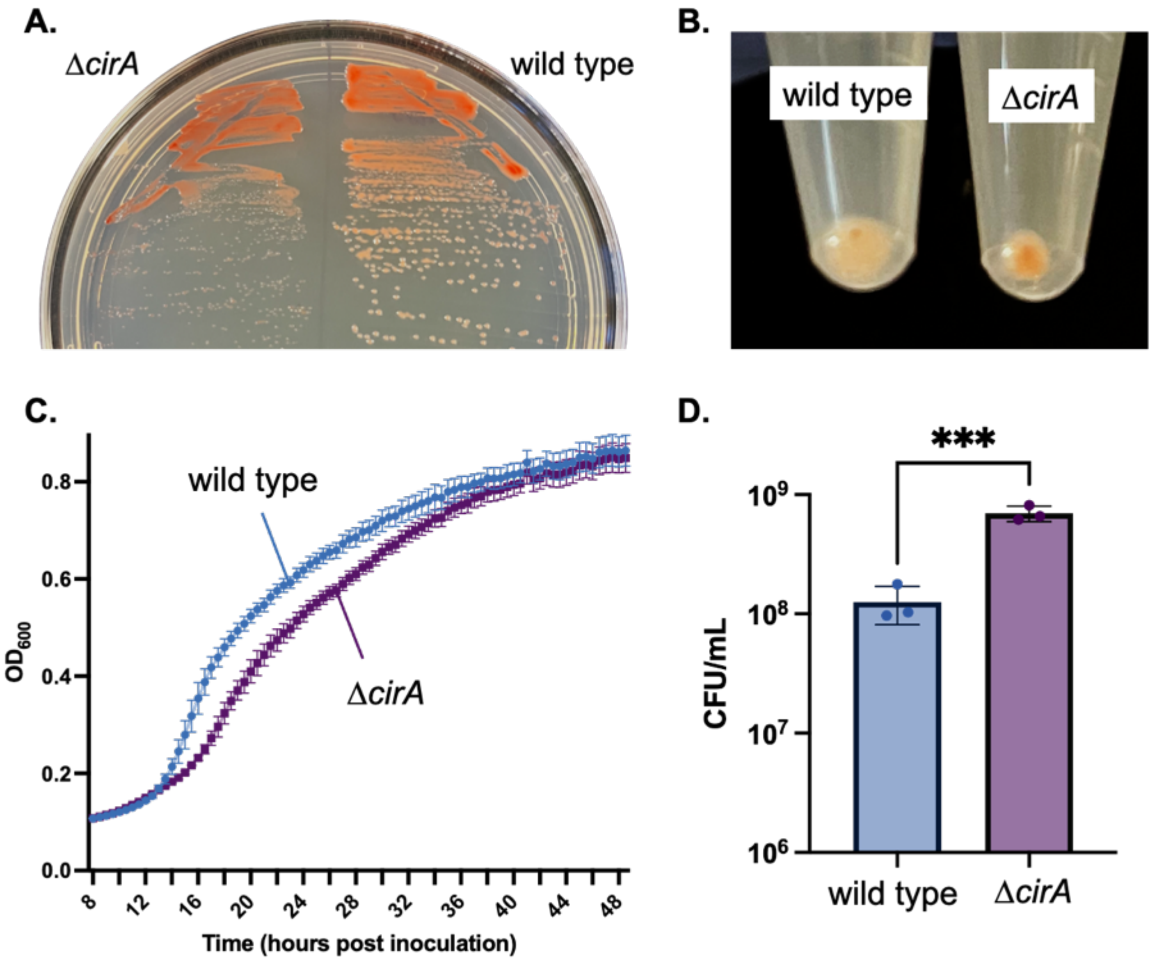


**Figures S3**

**Δ*cirA* colony morphology, pellet size, $OD_{600}$, and CFU count differ from the wild type. a)** Δ*cirA* and wild type *Hfx. volcanii* streaked on Hv-Cab solid agar plate (1.5% w/v). Plate incubated for 5 days at 45˚C before imaging. **b)** Cultures of *Hfx. volcanii* wild type and Δ*cirA* grown to $OD_{600}$ 0.55, and 1 mL of each culture was pelleted. **c)** Growth curve comparing wild type with Δ*cirA*. Cultures were inoculated overnight, then diluted to $OD_{600}$ 0.01. Eight replicates were inoculated into 96-well plate and $OD_{600}$ readings were taken at 30-minute intervals for two days. **d)** CFU/ml of wild type and Δ*cirA* strains at $OD_{600}$ of 0.5 in triplicate.
